# Adiponectin receptor agonist AdipoRon improves skeletal muscle function in aged mice

**DOI:** 10.1101/2021.09.16.460597

**Authors:** Priya Balasubramanian, Anne E. Schaar, Grace E. Gustafson, Alex B. Smith, Porsha R. Howell, Angela Greenman, Scott Baum, Ricki J. Colman, Dudley W. Lamming, Gary Diffee, Rozalyn M. Anderson

**Affiliations:** Department of Medicine, School of Medicine and Public Health, University of Wisconsin Madison, WI, United States; Department of Kinesiology, University of Wisconsin-Madison, Madison, WI, United States; Wisconsin National Primate Research Center, University of Wisconsin, Madison, WI, United States; Department of Cell and Regenerative Biology, University of Wisconsin, Madison, WI, United States; Geriatric Research, Education, and Clinical Center, William S. Middleton Memorial Veterans Hospital, Madison, United States

**Author notes:** Address for Correspondence Rozalyn M. Anderson, GRECC D5214, William S. Middleton Memorial Veterans Hospital, 2500 Overlook Terrace, Madison, WI 53705. Equal contributions (Priya Balasubramanian, Anne E. Schaar). Center for Geroscience and Healthy Brain Aging, University of Oklahoma Health Sciences Center, Oklahoma City, OK, United States.

**Keywords:** Muscle aging, mitochondria, PGC-1a, AMPK, adiponectin receptor

## Abstract

The loss of skeletal muscle function with age, known as sarcopenia, significantly reduces independence and quality of life and can have significant metabolic consequences. Although exercise is effective in treating sarcopenia it is not always a viable option clinically, and currently there are no pharmacological therapeutic interventions for sarcopenia. Here we show that chronic treatment with pan-adiponectin receptor agonist AdipoRon improved muscle function in male mice by a mechanism linked to skeletal muscle metabolism and tissue remodeling. In aged mice, 6 weeks of AdipoRon treatment improved skeletal muscle functional measures *in vivo* and *ex vivo*. Improvements were linked to changes in fiber type, including an enrichment of oxidative fibers, and an increase in mitochondrial activity. In young mice, 6 weeks of AdipoRon treatment improved contractile force and activated the energy sensing kinase AMPK and the mitochondrial regulator PGC-1a (peroxisome proliferator activated receptor gamma coactivator 1 alpha). In cultured cells, the AdipoRon induced stimulation of AMPK and PGC-1a was associated with increased mitochondrial membrane potential, reorganization of mitochondrial architecture, increased respiration, and increased ATP production. Furthermore, the ability of AdipoRon to stimulate AMPK and PGC1a was conserved in nonhuman primate cultured cells. These data show that AdipoRon is an effective agent for the prevention of sarcopenia in mice and indicate that its effects translate to primates, suggesting it may also be a suitable therapeutic for sarcopenia in clinical application.

## Introduction

Sarcopenia is the generalized and progressive decline in skeletal muscle mass that is accompanied by reduced muscle strength and reduced physical performance (Cruz-Jentoft and Sayer, 2019; Heymsfield et al., 2015; Santilli et al., 2014). Loss of skeletal muscle function significantly increases the risk for impaired mobility, falls, fractures, and morbidity (Angulo et al., 2016; Landi et al., 2013). Sarcopenia also has significant metabolic consequences; in humans, skeletal muscle is the largest insulin sensitive tissue in the body and its loss during aging dramatically increases the risk for sarcopenic obesity, diabetes, and cardiovascular disorders (Cleasby et al., 2016; Koster et al., 2011; Srikanthan et al., 2010; Stenholm et al., 2008). Exercise is the only known intervention to prevent sarcopenia (Joseph et al., 2012; Law et al., 2016; McLeod et al., 2019; Rebelo-Marques et al., 2018); however, mobility disability and polymorbidity can prevent its effectiveness as a means to counter skeletal muscle functional loss. Pharmacological therapeutic interventions to treat or prevent sarcopenia would be a considerable clinical advance but none have yet been identified.

Skeletal muscle composition is matched to function such that the muscles required for posture are not equivalent to those engaged in physical activity. Contractile content includes individual muscle fibers with differing force and endurance properties and differing metabolic fuel preference. Skeletal muscle also includes non-contractile components including satellite cells and stromal derived stem cells, vascular tissues, adipose, and extracellular matrix. With increasing age, muscle fibers undergo atrophy, and the levels of fibrosis and intra-muscular fat increase within and around the fiber bundles (Akazawa et al., 2017; Brioche et al., 2016; Lees et al., 2019; Mahdy, 2019; McGregor et al., 2014). Sarcopenia is associated with loss of mitochondrial activity (Andreux et al., 2018; Johnson et al., 2013; Petersen et al., 2015; Picca et al., 2018), although in a variety of species from rodents to nonhuman primates and humans it is the more oxidative fibers that are more resilient to age related atrophy (Crupi et al., 2018; Lexell, 1995; Murgia et al., 2017; Pugh et al., 2013). At the cellular level, aging is associated with insufficiency of fatty acid oxidation, and the accumulation of intracellular lipid (Choi et al., 2016; Koves et al., 2013; Koves et al., 2008). Importantly, our previous work in nonhuman primates show that age-related changes in mitochondrial energy metabolism, cellular redox, and lipid storage anticipate the onset of sarcopenia (Pugh *et al*., 2013), and occur in advance of physical activity decline and frailty (Yamada et al., 2013; Yamada et al., 2017). This suggests that changes in energy metabolism could be a contributing factor to age-related loss in muscle mass and function, and point to interventions that correct metabolic deficiency as potential agents for the treatment of sarcopenia.

Adiponectin is an adipose-derived peptide hormone that influences skeletal muscle cellular metabolism to activate lipid utilization and mitochondrial oxidative pathways (Shetty et al., 2012; Stern et al., 2016). Adiponectin signaling is transmitted via the ubiquitously expressed AdipoR1 and AdipoR2 receptors, with a poorly defined contribution from the T-cadherin receptor (Parker-Duffen et al., 2013; Yamauchi et al., 2007). A major effector protein in adiponectin receptor activation in skeletal muscle is the AMP activated protein kinase AMPK (Iwabu et al., 2010), a critical cellular energetic sensor (Burkewitz et al., 2014; Hardie et al., 2012). The actions of adiponectin somewhat resemble those of exercise: first, the adaptive response to exercise involves activation of metabolism in skeletal muscle and requires AMPK (Fentz et al., 2015; Lee-Young et al., 2009), and second, adiponectin and exercise share effectors downstream of AMPK including peroxisome proliferator-activated receptor gamma coactivator 1-alpha (PGC-1a), a key regulator of mitochondrial energy metabolism (Miller et al., 2019a) and a major player in the beneficial effects of exercise (Canto and Auwerx, 2010; Lira et al., 2010; Short et al., 2003). In fact, genetic approaches to increase expression of PGC-1a in skeletal muscle specifically enhance endurance and protect against muscle aging in mice (Benton et al., 2008; Ruas et al., 2012). AdipoR1 knockout studies reveal that adiponectin signaling is required for maintenance of skeletal muscle oxidative fibers in vivo and for exercise capacity, with both AMPK and PGC-1a implicated in its actions (Iwabu *et al*., 2010). AdipoRon is a small-molecule adiponectin receptor agonist, and treatment with AdipoRon corrects metabolic dysfunction in genetically obese and metabolically dysfunctional db/db mice (Okada-Iwabu et al., 2013). Although AdipoRon has been shown to impact gene expression in skeletal muscle in short term treatment of young mice, the ability of the drug to impact skeletal muscle in older mice is unknown.

## Results

### AdipoRon improves skeletal function and indices of insulin sensitivity in aged mice

To assess the ability of AdipoRon to influence skeletal muscle aging, AdipoRon (1.2mg/kg) or vehicle (PBS) were administered intravenously to 25-month old C57BL/6J male mice three times per week for 6 weeks. AdipoRon treatment resulted in a decline in body weight in aged animals which did not reach statistical significance (Fig.1A). A significant loss in fat mass measured by EchoMRI was observed in AdipoRon treated animals although there was no difference in lean mass compared to Controls (Fig.1B-C). Circulating levels of triglycerides were not different between Control and AdipoRon treated animals. In metabolic chamber assessments, food consumption, spontaneous activity, energy expenditure, or respiratory exchange ratio did not differ between Control and AdipoRon treated animals (Fig.S1). AdipoRon treated animals had significantly lower fasting glucose levels (Fig.1C) and a trend towards decreased fasting insulin levels (Fig.1D). In line with these findings, the HOMA-IR index of insulin resistance was significantly lower in AdipoRon treated animals (Fig.1E).

**Figure 1:**
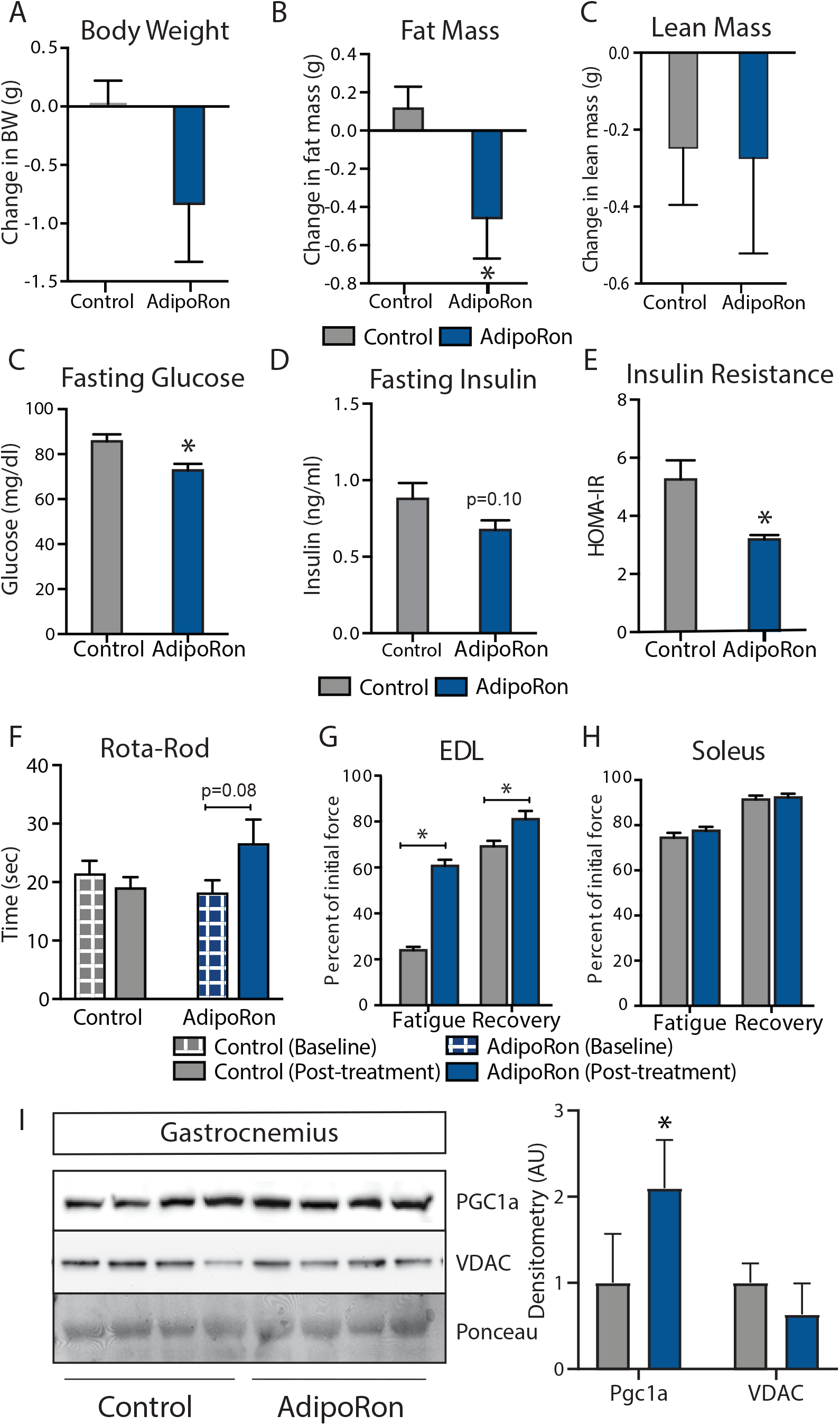
Metabolic and functional effects of chronic AdipoRon treatment in aged mice. Chronic AdipoRon treatment (1.2mg/kg BW, intravenous injection three times per week for 6 weeks) in aged male C57BL/6J mice (25 months old). (A) Change in body weight, (B) change in fat mass, (C) fasting glucose, (D) fasting insulin, (E) HOMA-IR. (F) *In vivo* latency to fall on Rota-Rod testing. Functional muscle changes assessed by measuring peak force after tetanic stimulation in *ex vivo* contractility experiments in isolated (G) extensor digitorum longus (EDL) and (H) soleus. (I) Immunodetection of mitochondrial marker VDAC and PGC1a protein expression in gastrocnemius. Control (n=7-10) and AdipoRon (n=6-8). Data shown as average ±SEM (*p<0.05).

The therapeutic potential of AdipoRon was evaluated via *in vivo* and *ex vivo* functional measures. Baseline measures for Rotarod did not differ between the groups; however, after 6 weeks of treatment, the AdipoRon treated animals tended to spend more time on the rotarod suggesting improved muscle function and agility (Fig.1F). *Ex vivo* contractility measurements in extensor digitorum longus (EDL, glycolytic) and soleus (oxidative) muscles were performed to determine the effects of AdipoRon treatment on whole muscle contractile behavior. Muscle maximal specific force generation and fatigability is known to be dependent on the type and quantity of expression of myosin heavy chains isoforms (Geiger et al., 2000). Consistent with this, the decline of tetanic force after initial simulation (fatigue) observed in Type II fiber enriched EDL was greater than that of Type I enriched soleus muscle in untreated aged mice. Force generation after repeated tetanus stimulation was significantly greater in the EDL from AdipoRon treated animals compared to controls, signifying a greater resistance to fatigue (Fig.1G). Electrical stimulation-induced force generation after a 20-minute recovery period was also significantly higher, indicative of greater muscle endurance following AdipoRon treatment. Interestingly, the beneficial effects of AdipoRon on fatigability were detected in EDL but not soleus muscle that had numerically greater values for both parameters at baseline compared to EDL (Fig.1H). These data suggest that oxidative muscles (soleus) are less sensitive to AdipoRon treatment compared to muscles with more reliance on glycolytic metabolism (EDL). PGC-1a is a key regulator of fiber type switching (Lin et al., 2002) and is engaged downstream of adiponectin receptor signaling (Iwabu *et al*., 2010). In gastrocnemius, a mixed fiber muscle in the lower leg with Type II fibers predominating, PGC-1a protein levels were higher from AdipoRon treated mice than from controls (Fig.1I), suggesting that activation of PGC-1a could be mediating the effects of AdipoRon in aged mice. Levels of VDAC (voltage dependent anion channel), an index of mitochondrial content, were not different between control and AdipoRon treated mice, indicating that mitochondrial biogenesis was not induced by AdipoRon in gastrocnemius.

### AdipoRon changes muscle composition including fiber-type specific metabolic activation

A change in muscle fiber type distribution within muscle is an outcome of chronic exercise (Mishra et al., 2015; Yan et al., 2011), where composition is adapted to functional demand. Differences in fiber type distribution within muscle is associated with distinct contractile properties (Zierath and Hawley, 2004). Gastrocnemius muscle was analyzed as representative of a mostly fast-glycolytic muscle (Type IIb and IIx muscle fibers). Immunohistological detection of myosin isoforms in gastrocnemius muscle cross-section (Fig.2A) revealed a significantly greater proportion of Type IIa myosin expressing fibers in muscle from AdipoRon treated mice compared to untreated control mice (Fig.2B). Histochemical detection of cytochrome c oxidase activity revealed an increase in intensity of COXIV activity staining in Type IIb fibers (Fig.2C). Interestingly, AdipoRon treatment was associated with a very small decrease in fiber size in Type IIb fibers (Fig.2D), indicating there may be structural adaptation to changes in mitochondrial activation status. Quantitative PCR confirmed Type IIb myosin isoform was by far the dominant fiber type in gastrocnemius (Fig.2E). AdipoRon treated mice had a modest reduction in expression of all myosin isoforms compared to controls although none of the differences were significant, and overall fiber type distribution was not altered (Fig.2E). Soleus muscle is a mostly slow-oxidative muscle, and RT-PCR confirmed that Type I and Type IIa myosin isoforms were dominant (Fig.S2). Although functional differences were not detected *ex vivo* in soleus, AdipoRon treatment significantly increased expression of Type I and Type IIx myosin, and numerically increased expression of Type IIa and Type IIb, although here too the overall distribution of fiber types was not altered (Fig.S2).

**Figure 2:**
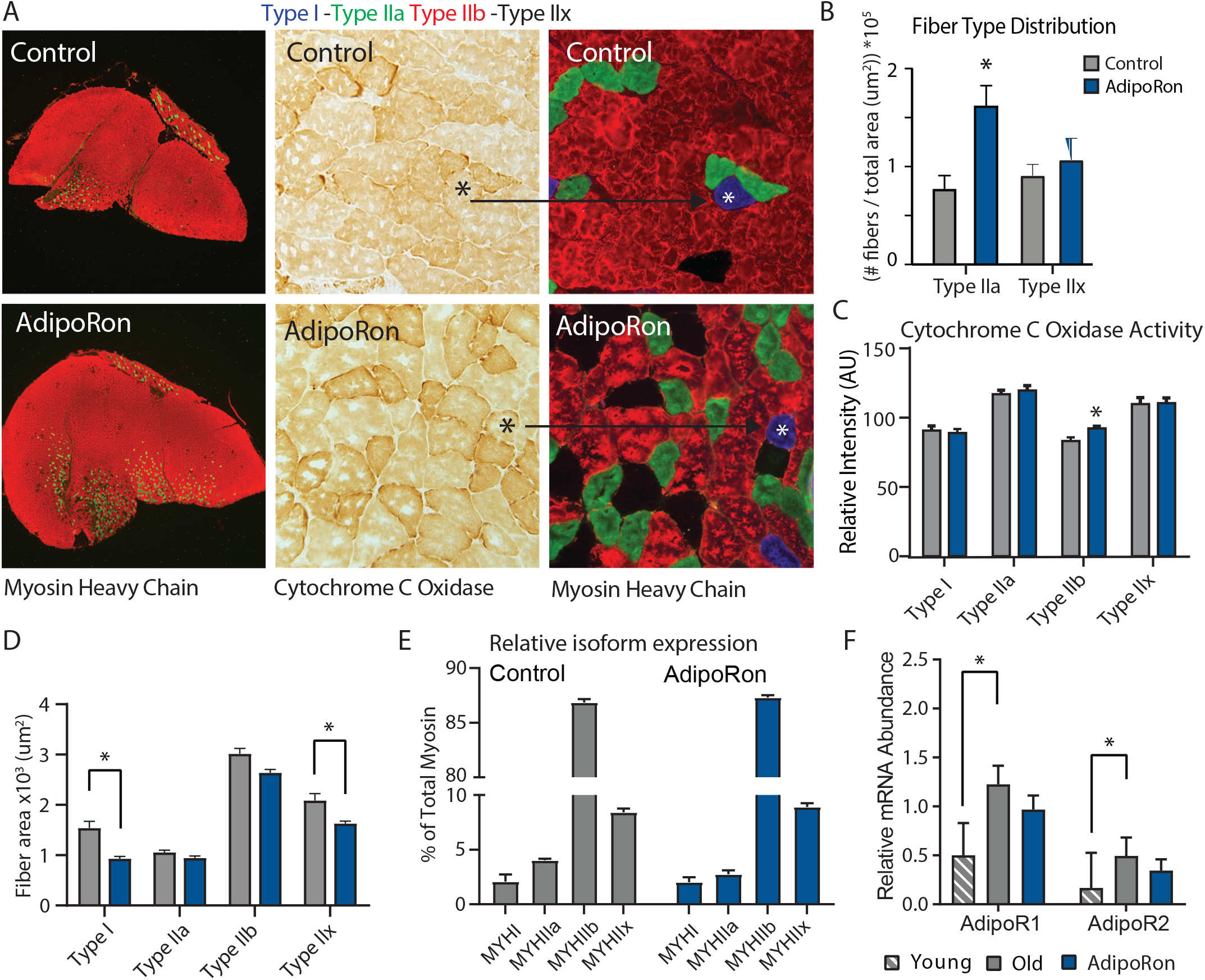
Fiber type specific metabolic and structural changes in response to AdipoRon. Chronic AdipoRon treatment in aged male C57BL/6J mice (25 months old). (A) Immunodetection of Type IIa (green) and Type IIb (red) shown at 2.5x magnification (Left), Representative histochemical staining for Cytochrome C Oxidase activity (Middle), and immunohistochemistry for Type I (blue), Type IIa (green), Type IIb (red), Type IIx (black) in adjacent gastrocnemius muscle sections at 20x magnification (Right). (B) Type IIa and IIx fiber counts normalized to (C) Cytochrome C Oxidase staining intensity analysis (D) and cross-sectional area of individual fibers. (E) Relative mRNA expression of myosin heavy chain isoforms. (F) Relative mRNA expression of adiponectin receptors AdipoR1 and AdipoR2 from young and aged control mice. Control (n=7-10) and AdipoRon (n=6-8). Data shown as log-fold change of mean dct (control – treatment) relative to 18s reference ± pooled SEM estimation and significance determined by Student t test (* p<0.05).

To investigate potential feedback from AdipoRon treatment, transcriptional expression of adiponectin receptors AdipoR1 and AdipoR2 was determined. Levels of AdipoR1 were significantly greater than AdipoR2 in gastrocnemius (Fig.2F). Levels of both receptors were higher in gastrocnemius of old mice compared to young mice, corroborating previous reports that adiponectin receptor expression changes with age (Ito et al., 2018). Expression levels of both receptors were numerically lower in AdipoRon treated mice. In soleus muscle, AdipoR1 expression was not altered with age (Fig.S2) and AdipoRon increased expression, although neither change reached significance. AdipoR2 levels declined with age in soleus of control mice, but not in mice treated with AdipoRon. The known age-associated increase in fibrosis in muscle (Wood et al., 2014) prompted an investigation of the effects of AdipoRon treatment on collagen content in aged animals. Histochemical staining for collagen showed that over this relatively short treatment period AdipoRon did not impact collagen content or fibrosis in the aged muscle (Fig.S3). Together, these data show that chronic AdipoRon treatment mimics exercise including changes in fiber type composition with more oxidative fibers, a shift in glycolytic muscle fibers towards a more oxidative phenotype, and a correction of age-related elevation of adiponectin receptor expression.

### AdipoRon improves contractile force in young mice in a muscle-type-specific manner

To assess the chronic effects of AdipoRon on metabolism and muscle function in young mice, we administered AdipoRon (1.2mg/kg) or vehicle (PBS) three times per week for 6 weeks by tail vein injections beginning at 2 months of age in male B6C3F1 hybrid mice. Chronic AdipoRon treatment did not alter body weight (Fig.3A) or fasting glucose levels in young animals (Fig.3B). *Ex vivo* contractility measurements were performed in EDL and soleus muscles from the young Control and AdipoRon treated mice. Chronic AdipoRon treatment resulted in significantly improved ability to recover after repeated tetanus stimulation in EDL muscle suggesting higher resistance to fatigue (Fig. 3C). Consistent with results from the aged mice, the effect of AdipoRon was muscle-type specific as it did not impact measures of contractility and force in the soleus muscle (Fig.3D). This confirms that AdipoRon preferentially improves function in fast muscle with predominantly Type IIb/IIx fibers independent of the age of the mice during treatment. PGC-1a protein levels were higher in gastrocnemius from AdipoRon treated mice than from controls (Fig.3E). Total transcript levels of PGC1a were higher in gastrocnemius from AdipoRon treated mice but not in soleus (Fig.3F), although transcript levels of selected PGC1a target genes (*Cpt1a, Acox1* and *Acadm*) were not altered by AdipoRon treatment (Fig.3G). The *Ppargc1* gene produces several isoforms including PGC-1a1 and PGC-1a4 from the canonical promoter and variants PGC-1a2, and PGC-1a3 from an alternate promoter (Ruas *et al*., 2012). AdipoRon altered the profile of PGC-1a isoforms indicating a change in promoter preference (Fig. 3H).

**Figure 3:**
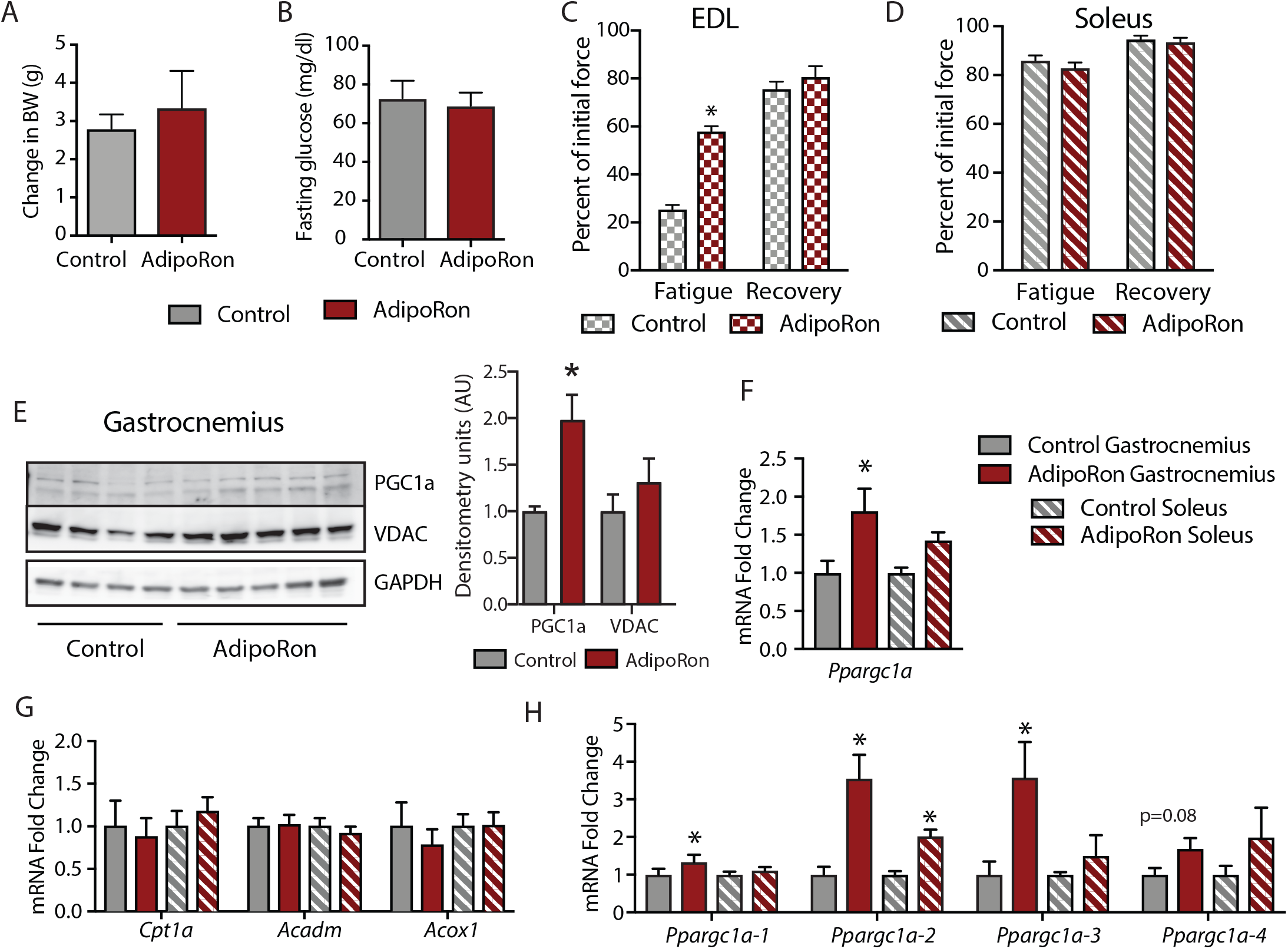
Metabolic and functional effects of chronic AdipoRon treatment in young mice. Chronic AdipoRon treatment (1.2mg/kg BW, intravenous injection three times per week for 6 weeks) in young male B6C3F1 mice (3.5 months). Measures of (A) Body weight, (B) fasting glucose in Control (n=5) and AdipoRon (n=5) treated mice. Measures of peak force after tetanic stimulation in ex vivo contractility experiments in isolated EDL (C) and Soleus (D) muscles. (E) Immunodetection of protein levels of PGC1a and VDAC in gastrocnemius muscle. (F) mRNA expression of Ppargc1a, PGC-1a gene targets, and Ppargc1a isoforms in gastrocnemius and soleus muscle. Control (n=10) and AdipoRon (n=8) animals. Data shown as average shown as average ±SEM and significant difference between group means determined by Student’s t-test (* p<0.05).

To determine whether AdipoRon mediated changes in muscle function were coincident with changes in muscle composition, tissue sections from young mice were subject to immunodetection of myosin heavy chain isoforms. No changes in fiber cross sectional area were identified, presumably because young mice are in peak physical condition (Fig.S2). Histochemical detection of maximal cytochrome c oxidase (Complex IV, COXIV) and succinate dehydrogenase (Complex II, SDH) activity (Mahad et al., 2009), similarly showed no differences in young mice with or without AdipoRon treatment (Fig.S2). Fiber area measurements revealed that the fiber size was inversely proportional to mitochondrial activity (TypeIIb>IIx>I>IIa), which has been described previously as the muscle fiber type-fiber size paradox (van Wessel et al., 2010).

### Acute treatment with AdipoRon activates PGC-1a in a muscle type-specific manner

To understand the differences among muscle groups in terms of PGC-1a expression and adiponectin receptor distribution, gene expression analysis was conducted in extracts from gastrocnemius and soleus muscles from young male B6C3F1 hybrid mice (6 weeks old). Endogenous levels of *Ppargc1a* expression were significantly higher in soleus than gastrocnemius muscle (Fig.4A), which is to be expected due to the higher dependence of soleus muscle on oxidative metabolism (Koves et al., 2005). Interestingly, the distribution of isoforms was largely equivalent between the two muscle types, with *Ppargc1a1* being the dominant isoform, lesser contributions from *Ppgargc-1a4* and *Ppargc1a2*, and lowest levels detected for *Ppargc1-a3* (Fig.4B). The possibility that differences in responsivity to AdipoRon between the gastrocnemius and soleus muscle types observed in the chronic treatment studies could be due to differences in adiponectin receptor expression was investigated. Gene expression analysis revealed that in young mice there was no overt difference in the expression levels of *AdipoR1* and *AdipoR2* between gastrocnemius and soleus, though *AdipoR1* expression was notably higher than AdipoR2 for both muscle types (Fig.4C).

**Figure 4:**
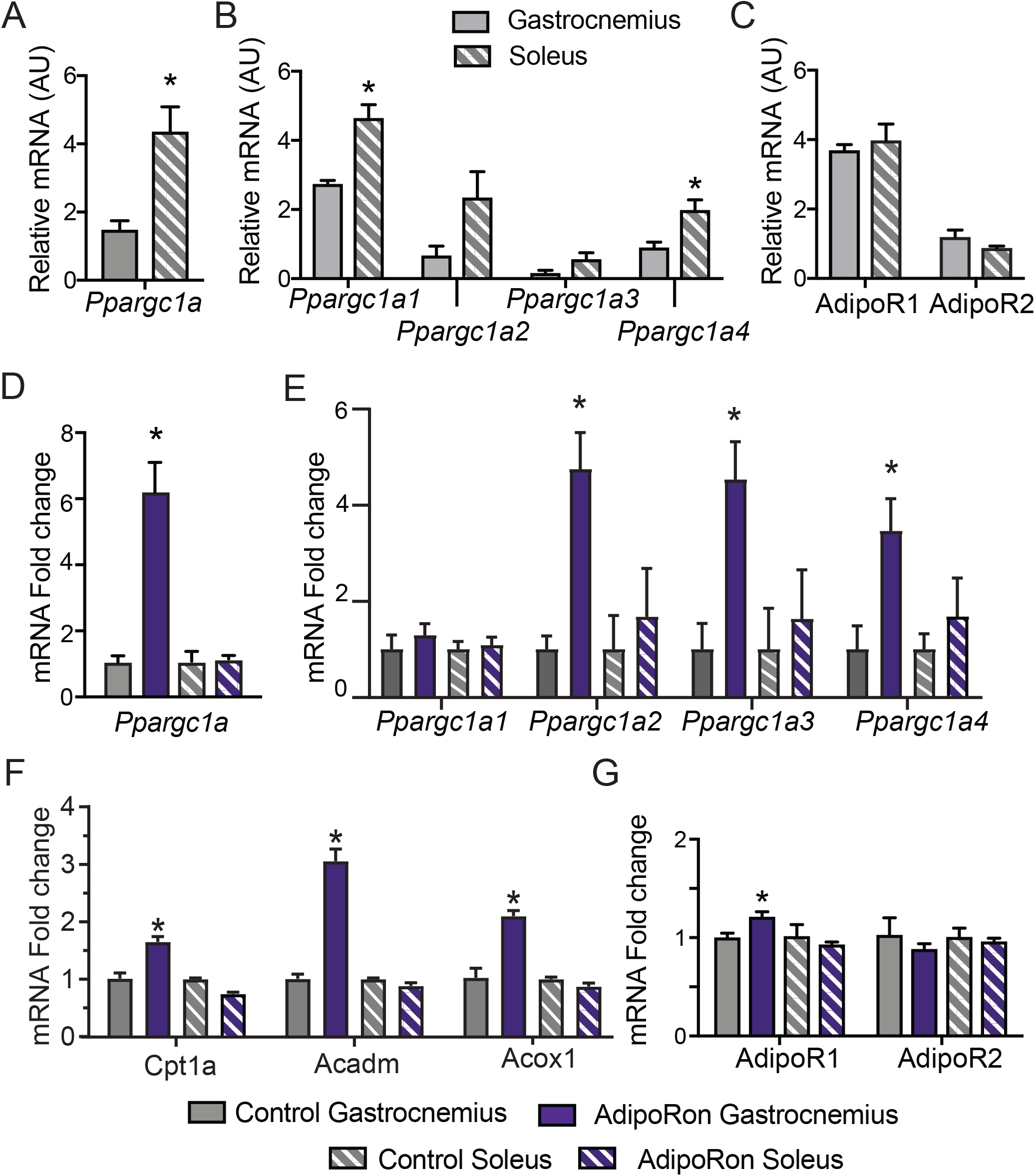
Muscle specific transcriptional response to acute AdipoRon treatment. Comparison of relative mRNA expression between gastrocnemius and soleus muscles of (A) Ppargc1a, (B) Ppargc1a isoforms, and (C) AdipoR1and R2 in muscles from Control mice. Changes in gene expression of (D) Ppargc1a, (E) Ppargc1a isoforms, (F) gene targets of PGC1a and (G) gene expression of adiponectin receptors R1 and R2 in gastrocnemius and soleus muscle of 6 week old male B6C3F1 hybrid mice 90-minutes post-IV injection with AdipoRon (1.2mg/kg BW), Control (n=3) and AdipoRon (n=5). Data shown as average shown as average ±SEM and significant difference between group means determined by Student’s t-test (* p<0.05).

To investigate the early response to AdipoRon in skeletal muscle, the impact of a single intravenous injection of AdipoRon (1.2mg/kg BW) was investigated in muscles from young male B6C3F1 hybrid mice (6 weeks old) collected 90 minutes post-injection. AdipoRon treatment increased the gene expression of *Ppargc1a* in gastrocnemius but not soleus muscle (Fig.4D). Exercise training has been reported to induce the expression of the Canonical promoter driven *Ppargc1a1* and *Ppargc1a4*, with a lesser effect on isoforms expressed from the Alternate promoter. Acute treatment with AdipoRon resulted in an upward trend of transcript levels of most highly expressed isoform *Ppargc1a1*, a significant increase in the of *Ppargc1a4* isoform in gastrocnemius muscle, although increases in expression of *Ppargc1a2* and *Ppargc1a3* did not reach significance (Fig. 4E). No changes were seen with the expression of *Ppargc1* isoforms in soleus muscle, and gene targets of PGC-1a involved in fatty acid metabolism (*Cpt1a, Acox1* and *Acadm*) were increased in response to acute AdipoRon treatment in gastrocnemius but not in soleus muscle (Fig. 4F). Exercise training has also been linked to changes in adiponectin receptor expression where an 8-week exercise regimen increased bicep femoris muscle mRNA expression of *AdipoR1* but had no effect of *AdipoR2* (Huang et al., 2006). Within our study, AdipoRon similarly increased the expression of *AdipoR1* in gastrocnemius muscle but had no effect on levels of *AdipoR2* (Fig.4G). AdipoRon had no effect on the expression levels of adiponectin receptors in the soleus muscle, confirming that the effects of AdipoRon are muscle-type specific.

### AdipoRon stimulates mitochondrial function in cultured fibroblasts

Adiponectin receptor stimulation by AdipoRon has been linked to activation of AMPK and PPARa (peroxisome proliferator activated receptor alpha) pathways in muscle and liver, including increased PGC-1a transcription and activation of expression of its gene targets (Okada-Iwabu *et al*., 2013). Studies of renal function in a genetic model of diabetes (db/db) show AMPK and PPARa engagement by AdipoRon and have revealed changes in lipid parameters in response to AdipoRon in renal endothelial cells (Choi et al., 2018). Less is known of the functional consequence of PGC-1a activation by AdipoRon as it relates to cellular metabolism and mitochondrial function. Cultured NIH-3T3 fibroblasts were treated for 48 hrs with 50µm AdipoRon treatment. Immunodetection of PGC1a protein and VDAC (Voltage dependent anion channel) revealed a significant increase in PGC-1a protein and a more modest but significant increase in VDAC, an index of mitochondrial content (Fig.5A). Mitochondrial membrane potential measured by JC-1 assay was increased after 2 and 4 hrs of 50µm AdipoRon treatment (Fig.5B). The impact of AdipoRon on mitochondrial function was coincident with an increase in cellular oxygen consumption measured over the same time period (Fig.5C). Cellular ATP concentration was significantly higher in AdipoRon treated fibroblasts 12 hours following 50µm treatment (Fig.5D). Our previous studies have shown that increased respiration is accompanied by slower growth in cultured fibroblasts (Miller et al., 2019b). Consistent with this, cell proliferation was negatively impacted by 24 hrs exposure to AdipoRon regardless of seeding density (Fig.5E). We had previously shown that mitochondrial organization is influenced by chronic elevation of PGC-1a (Miller *et al*., 2019b). Mitochondria were visualized in fixed fibroblasts by immunofluorescence detection of TOMM20 following 24 hrs AdipoRon treatment (Fig.5F). Treated cells showed quantitatively significant differences in fluorescent intensity and in mitochondrial network architecture as mitochondria tended toward shorter branch morphologies. These data reveal prominent effects of acute AdipoRon treatment on the mitochondrial regulator PGC-1a and on mitochondrial function and organization and link these PGC-1a-associated metabolic changes to delayed proliferation and growth.

**Figure 5:**
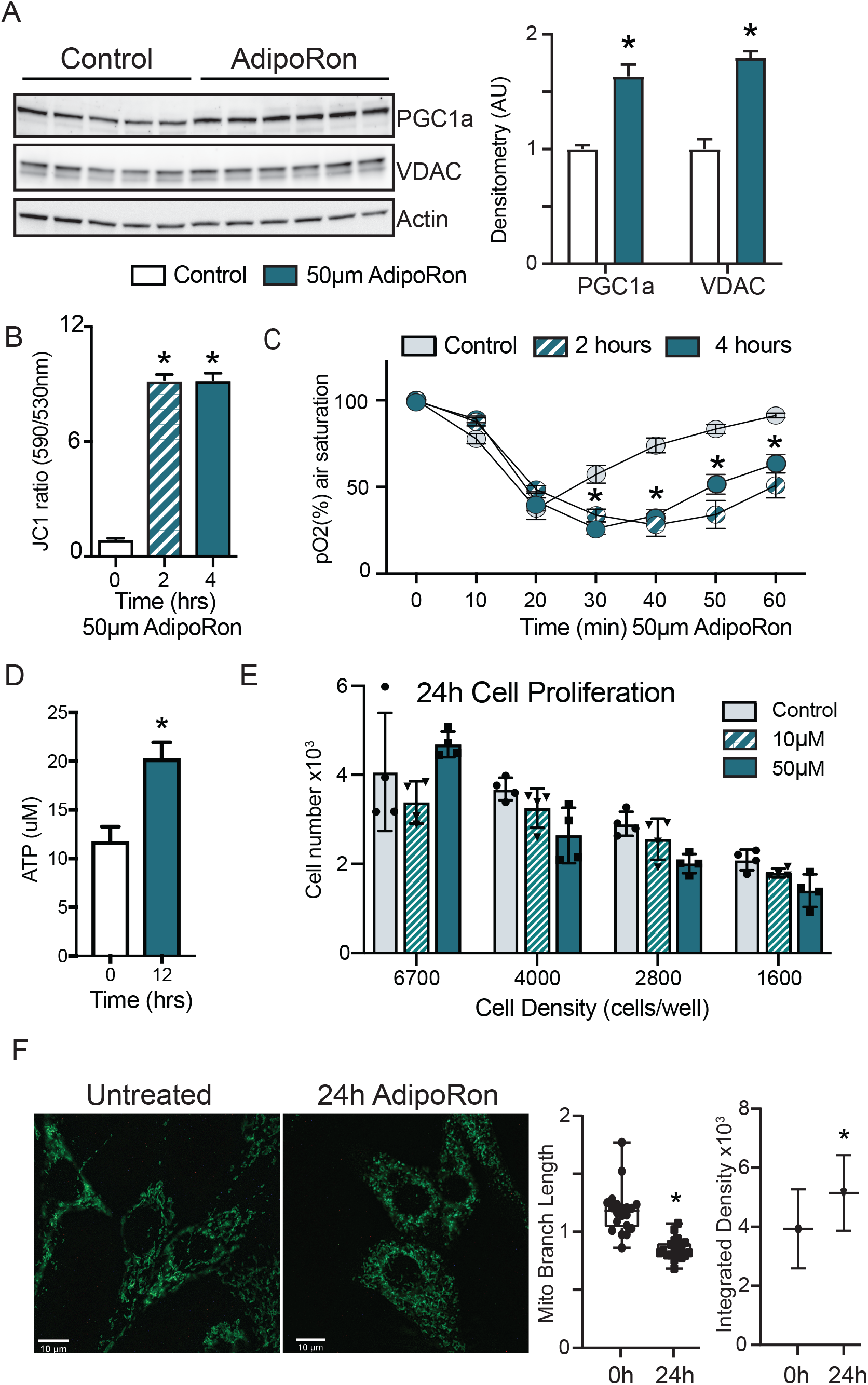
Short term impact of AdipoRon treatment on mitochondrial metabolism in murine fibroblasts. (A) Western blot showing protein levels of PGC1a and VDAC after 48 hrs of 50µm AdipoRon treatment of NIH-3T3 fibroblasts (n=5-6 per group), (B) Mitochondrial membrane potential measured by JC-1 assay at 2 and 4 hrs of AdipoRon treatment (n=6 per group), (C) Oxygen consumption measured in response to AdipoRon treatment (Control-open circle, AdipoRon 2hrs-hatched circle, AdipoRon 4hrs-filled circle) by oxoplate assay (n=5 per group), (D) ATP concentration after indicated time with 50µm AdipoRon treatment, (E) Impact of 24 hrs of AdipoRon treatment on cell proliferation of differing seeding densities as assessed by CyQUANT assay (n=4 per group). (F) Fluorescent detection of mitochondria in fixed NIH-3T3 following 48 hours AdipoRon treatment including quantitation of integrated density (product of mean intensity and mitochondrial area), mitochondrial footprint, and mean mitochondrial skeleton branch length. N=15-20 cells per timepoint. Data shown as average shown as average ±SEM (* p<0.05 via Student’s t-test).

### AdipoRon actions are conserved in nonhuman primate cultured cells

In cultured muscle cells AdipoRon activates AMPK and increases the expression of PGC1a and its gene targets (Ito *et al*., 2018; Okada-Iwabu *et al*., 2013). The degree to which AdipoRon action is conserved among other cell types and among species remains an open question. Our data revealed functional changes in mitochondria in response to AdipoRon treatment in NIH3T3 fibroblasts and we sought to investigate whether the molecular underpinnings for these observations were linked to AMPK and to PGC-1a. NIH3T3 fibroblasts were treated with AdipoRon treatment (10 or 50µm) for 10 min. Increased levels of activating phosphorylation of AMPK at Thr172 were detected in extracts of AdipoRon treated cells compared to controls for both the lower and higher dose (Fig.6A). These data align with prior reports of AdipoRon directed AMPK activation in C2C12 myoblasts and glomerular endothelial cells (Choi *et al*., 2018; Ito *et al*., 2018; Okada-Iwabu *et al*., 2013). Increased levels of *Ppargc1a* transcript were detected after 90 min of AdipoRon treatment at 50µm but not 10µm concentration (Fig.6B). Although the transcript of PGC-1a itself was not increased at the lower dose, expression levels of known targets of PGC-1a (*Cpt1a, Acox1* and *Acadm*) were significantly increased at 10µm and at 50µm, indicating that PGC-1a activation at the lower dose of AdipoRon may occur at the post-transcriptional level. The response to AdipoRon was next investigated in a diverse cell population derived from nonhuman primates. Peripheral blood mononuclear cells (PBMCs) were isolated from blood taken from adult male rhesus monkeys. The cells were cultured overnight and subsequently treated with AdipoRon. Activating phosphorylation of AMPK at Thr172 was increased after 10 min of AdipoRon treatment at 50µm but not 10µm concentration (Fig.6C). One hour following AdipoRon treatment of PBMCs significant increases in transcript levels of *Ppargc-1a* and PGC-1a gene targets were detected at both the lower and higher doses (Fig.6D). These data demonstrate that the molecular response to AdipoRon is conserved among cell types and indicate that AdipoRon may be effective in nonhuman primates.

**Figure 6:**
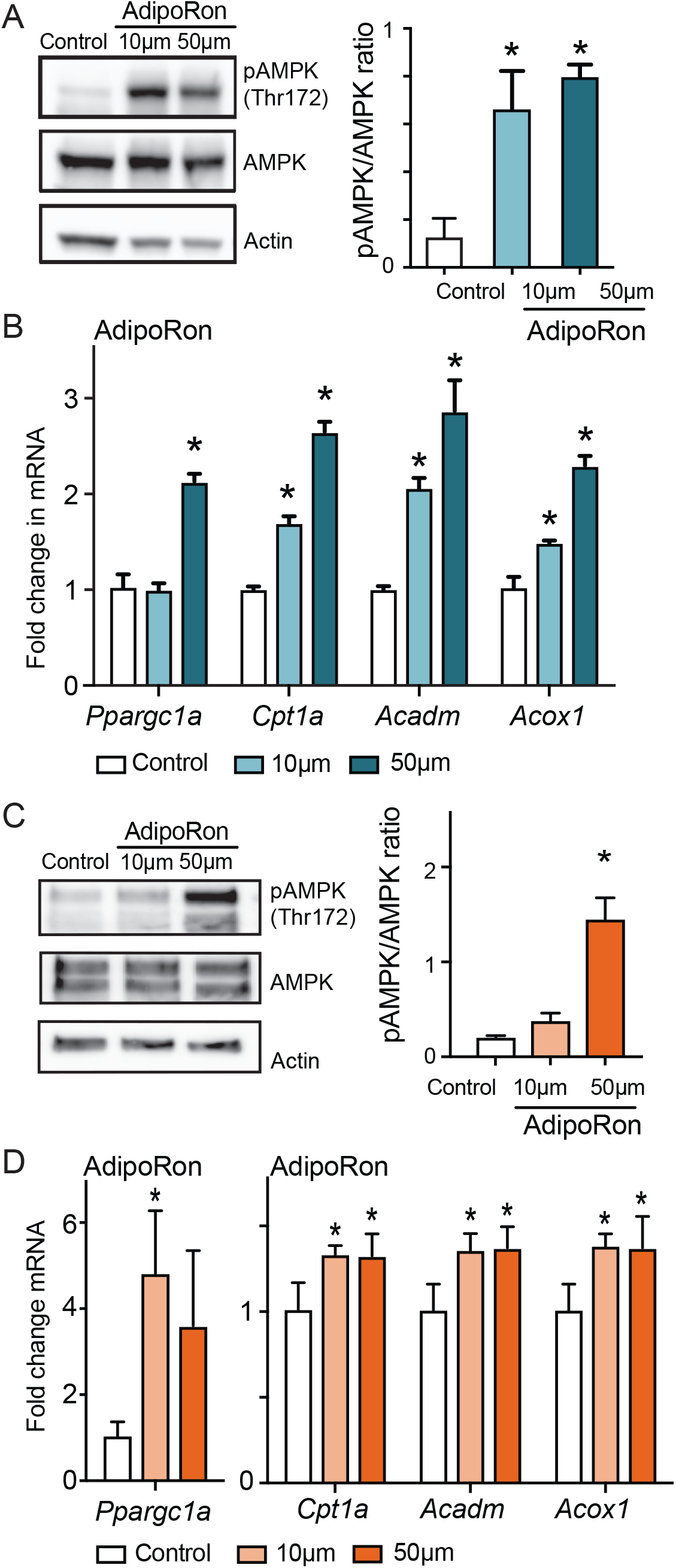
AdipoRon activates AMPK, increases the expression of PGC1a and its gene targets in nonhuman primate cultured cells. (A) Phosphorylation of AMPK at Thr172 after AdipoRon treatment (10 or 50µm) for 10 minutes (n=3/group) in NIH-3T3 fibroblasts, (B) Transcript levels of PGC1a and its gene targets after 90 minutes of AdipoRon treatment (10 or 50µm, n=4-5/group) (C) Phosphorylation of AMPK at Thr172 after AdipoRon treatment for 10 minutes in cultured nonhuman primate peripheral blood mononuclear cells (PBMCs) (n=3 per group) (D) Transcript levels of PGC1a and PGC1a gene targets after 60 minutes of AdipoRon treatment (n=3-4 per group). Data shown as average ±SEM (* p<0.05).

## Discussion

We have provided evidence of the therapeutic potential of AdipoRon for the treatment of sarcopenia in mice. *In vivo* and *ex vivo* functional assessments of skeletal muscle revealed that 6 weeks of AdipoRon treatment improved skeletal muscle function in aged mice. Interestingly, the effect was muscle type specific, where AdipoRon impacted functional, metabolic, and gene expression outcomes in glycolytic muscles (EDL and gastrocnemius) but not in highly oxidative soleus muscle. Skeletal muscle is known for its heterogeneous population of muscle fibers, each with distinct structural, biochemical, and metabolic properties. Studies in humans and non-human primates have previously demonstrated that there is a selective decrease in fiber size and mitochondrial activity of Type II fibers with aging (Brunner et al., 2007; Murgia *et al*., 2017; Pugh *et al*., 2013). AdipoRon treatment increased mitochondrial activity in a fiber-type specific manner in the gastrocnemius muscle of aged mice and resulted in an increase in the number of Type IIa fibers. Given the narrow time frame of the intervention and prior links between PGC-1a expression and fiber type switching (Lin *et al*., 2002), it seems more likely that AdipoRon remodels existing fibers rather recruiting new fibers from the satellite pool. *In situ* histochemical staining for cytochrome c oxidase activity combined with fiber typing in muscle cryosections allowed us to investigate the structural and metabolic effects of AdipoRon based on fiber types. One of the striking findings of our study is that AdipoRon distinctively augmented mitochondrial metabolism in Type IIb fibers in aged mice. At the whole tissue level, this effect translated to increases in the force of contractility and endurance in the EDL, a glycolytic muscle that is rich in Type IIb fibers but not in the soleus muscle that is rich in Type I fibers.

Consistent with previously published studies (Okada-Iwabu *et al*., 2013), AdipoRon had a positive impact on glucoregulatory function in aged mice. In young, fit mice the effects of AdipoRon at the cellular level were conserved but the systemic glucoregulatory and functional outcomes were not. It seems likely that AdipoRon may not be effective to stimulate muscle function outside of an existing age-related, genetic, or diet induced decline. Experiments in isolated muscle preparations may shed light on the tissue autonomous versus systemic effects of AdipoRon on skeletal muscle metabolism, particularly in the context of aging. In our study, the beneficial impact of AdipoRon treatment in skeletal muscle was mechanistically linked with PGC1a activation. Previous studies on muscle-specific overexpression of PGC1a have reported similar findings, however in those genetic models, whole body insulin sensitivity was adversely affected (Benton *et al*., 2008; Choi et al., 2008; Handschin et al., 2007; Wong et al., 2015). Our results show that AdipoRon treatment resulted in increases in expression of endogenous *Ppargc1a* mRNA with concomitant modest but significant increases in PGC-1a protein levels. These data suggest that the activation of regulatory pathways upstream of PGC-1a may be key to exerting beneficial effects beyond skeletal muscle energetics, although we cannot rule out systemic effects of AdipoRon to alter metabolism in other tissues playing a role in glucoregulatory outcomes.

A decline in muscle energy metabolism precedes the onset of sarcopenia in nonhuman primates (Pugh *et al*., 2013), a finding that promoted this study of AdipoRon as a potential therapeutic agent. Our data show not only the activation of AMPK and of PGC-1a, but also reveal functional metabolic changes in response to AdipoRon treatment, in tissues as described above but also in cultured cells. In NIH3T3 fibroblasts mitochondrial membrane potential, cellular respiration, proliferation, and mitochondrial architecture were all impacted by AdipoRon. The inference is a shift toward fatty acid fuel utilization, aligning with prior reports of lipid remodeling outcomes reported in renal endothelial cells (Choi *et al*., 2018). These outcomes are also consistent with our genetic studies of modest PGC-1a overexpression in preadipocytes, where these same metabolic changes induced in mitochondria were linked to greater capacity for lipid fuel utilization, changes in lipid handling, lipid storage, and membrane lipid composition (Miller *et al*., 2019b). Anti-inflammatory actions of AdipoRon could also exert beneficial muscle outcomes as recently reported in a genetic mice model of Duchenne muscular dystrophy and warrants investigation in our aging study (Abou-Samra et al., 2020). The fact that AdipoRon is equally potent in activating AMPK and PGC1a in PBMC’s demonstrates that the findings are likely translatable to nonhuman primates and perhaps humans. In addition, activation of adiponectin signaling has implications for aging and longevity as it shares signaling mechanisms involving AMPK and PGC1a with a known anti-aging intervention, calorie restriction (CR) (Balasubramanian et al., 2017). CR is associated with higher circulating adiponectin, especially the high molecular weight (HMW) isoform, which is the most potent in insulin sensitization (Miller et al., 2017). One limitation of that study, shared in this study, is that the work involved only male mice. Our ongoing studies in mice and in nonhuman primates point to profound sex dimorphism in function, tissue level, and cellular effects of muscle aging. It will be critical to extend these studies of AdipoRon as a potential intervention to prevent and treat sarcopenia to female mice, and in translating the work to nonhuman primates to include both male and female monkeys. Our data suggest that AdipoRon, through its actions on mitochondrial metabolism, skeletal muscle composition, fiber fatigability and function, and physical capacity, is a promising therapeutic agent to preserve skeletal muscle function in aged populations.

## Materials and Methods

### Mice and Treatment

Two cohorts of mice were used. Six-week-old male B6C3F1 hybrid mice were obtained from Jackson Laboratories (Bar Harbor, ME, USA) and 24-month old male C57BL/6J mice from the NIA aged mouse colony. The animals were housed under controlled pathogen-free conditions. The animals were fed ad-libitum Purina LabChow 5001 diet with free access to water, and placed on a 12h light/dark cycle. AdipoRon (10mg) (Adipogen, CA, USA) was reconstituted with 500ul of DMSO (stock) and stored at -20C. For the acute experiment, the animals were given either a single intravenous dose of AdipoRon @1.2 mg/kg BW in PBS (n=5) or an equivalent volume of DMSO in PBS (n=3, Controls) in a total volume of 100ul. Extensor digitorum longus (EDL) and soleus muscle groups were harvested after 90 minutes. For the chronic experiments in young B6C3F1 mice (n=12) and old C57BL/6J mice (n=20), the animals in each age group were divided into 2 groups and received either AdipoRon @1.2 mg/kg BW in PBS or an equivalent volume of DMSO in PBS via tail vein injections three times per week (Monday, Wednesday and Friday) for 6 weeks. Body weight and food intake measurements were recorded once a week. Body composition was measured at the beginning and at the end of the treatment period using EchoMRI Body Composition Analyzer (Houston, TX). At the end of the treatment period, animals were sacrificed. EDL, soleus, and gastrocnemius muscle specimens were mounted in OCT oriented to present fiber cross-section for tissue sectioning or snap frozen in liquid nitrogen and stored at -80°C. All animal protocols were approved by the Institutional Animal Care and Use Committee at the University of Wisconsin, Madison.

### Fasting glucose and insulin measurements

Overnight fasting blood samples were collected for measures of glucose (One touch Ultra Blue glucometer), and insulin levels (Ultra-Sensitive Mouse Insulin ELISA Kit (CystalChem, IL, USA)). HOMA-IR (homeostasis model assessment of insulin resistance) index was calculated as [fasting serum glucose × fasting serum insulin/22.5].

### Metabolic Chambers

Metabolic parameters and activity were measured (Columbus Instruments Oxymax/CLAMS metabolic chamber system (Columbus, OH)) in mice acclimated to housing in the chamber for approximately 24 hours followed by 24hr continuous data collection period.

### Ex vivo muscle force measurements

EDL and soleus hindlimb muscles were dissected and perfused with oxygenated (95% O_2_, 5% CO_2_) Tyrode’s solution at room temperature (NaCl 145 mM, KCl 5 mM, CaCl_2_ 2mM, MgCl_2_ 0.5 mM, NaH_2_PO4 0.4 mM, NaHCO_3_ 24 mM, EDTA 0.1 mM, Glucose 10 mM). Isolated muscles were attached to a contractile apparatus capable of measuring force (Aurora Scientific) and were electrically stimulated using parallel platinum electrodes. Maximal twitch force and tetanic force were determined by adjusting the length of the muscle until maximal twitch force is reached, defining the optimal length. A 15-minute period of rest allowed muscle to equilibrate to the new environment. Fatigability was defined as the decline in tetanic force following 10 minutes of continuous stimulation. Muscles were tetanically stimulated at 100 Hz for 500 ms every 5 seconds at a voltage that generates the maximal force. Following 10 minutes of fatiguing stimulation, a 20-minute recovery period was allowed, where there was no stimulation in an oxygenated Tyrode’s perfusion. After the recovery period, muscle was electrically stimulated again to quantify the amount of recovery force. A recovery force higher relative to fatiguing force was taken as an indication that the decline in force during fatiguing stimulation is reversible. The amount of fatigue and recovery force was represented as a percentage of the initial force (before fatiguing stimulation). Following these measurements, fatigued muscles were weighed, quick frozen in liquid nitrogen and stored at -80°C.

### Western blotting

Equal amounts of skeletal muscle protein extract (45µg) were separated on Mini-Protean TGX precast protein gels (Biorad, CA, USA) and transferred to a PVDF membrane using a Trans-Blot semi-dry transfer system (Biorad, CA, USA). The membranes were blocked using 5% BSA in TBST for phospho-antibodies or 5% non-fat milk for other antibodies for 1hr at RT. The following primary antibodies were used overnight: pAMPK, AMPK and GAPDH (Cell signaling, Beverly, MA, USA), PGC1a (H300, Santa Cruz) and Beta-Actin (Sigma), washed in TBST, and incubated with the respective HRP-conjugated secondary antibodies (Vector Labs, Burlingame, CA, USA) for 1 hr at RT. Proteins were detected (Supersignal West Pico or Femto Chemiluminescent substrate solutions (Thermofisher Scientific)) and digital images acquired (GE ImageQuant Gel Doc Imaging system). Densitometric analysis was carried out using Fiji software. **Real time PCR:** RNA was extracted in Trizol reagent and isolated (Direct-zol RNA Miniprep kit (ZymoResearch, Irvine, CA, USA)). cDNA was synthesized from 1µg of RNA (High-Capacity cDNA Reverse Transcription Kit (Applied Biosystems)). Real time PCR reactions were carried out using iTaq Universal SYBR Green mix (Biorad) with 10ng cDNA per reaction. The primer sequences are listed in Table S1 and the data was analyzed by 2^^^ct^ method (Livak and Schmittgen, 2001).

### Histology

Serial OCT mounted cryostat sections (10 µm in thickness) were cut at -14°C (Leica Cryostat (Fisher Supply, Waltham, MA)). Freshly cut sections were stained for cytochrome c oxidase activity and digital image captured the same day as described previously (Pugh *et al*., 2013). Briefly, sections were air-dried at RT for 10-15 minutes, incubated in a solution of 0.1M phosphate buffer, pH 7.6, 0.5 mg/ml DAB (3,3’-diaminobenzidine), 1 mg/ml cytochrome c, and 2 µg/ml catalase at RT for ∼30 minutes, washed with PBS, dehydrated, cleared and mounted under a glass coverslip (Permount (Fisher)). Similarly for succinate dehydrogenase activity staining, the frozen muscle sections were air-dried for 10-15 minutes and incubated in SDH reaction mixture containing [1.5mM nitroblue tetrazolium (NBT) and 50mM di-sodium succinate in 0.2M PBS (pH 7.6) at RT for ∼20 minutes, washed with PBS, dehydrated, cleared and mounted under a coverslip with Permount (Punsoni et al., 2017). Muscle fibrosis was determined by trichrome staining (Abcam, MA). For stain intensity and/or stain area analysis in digital images (n=5 per tissue), see below.

### Immunofluorescence

Fiber types (I, IIa, IIb and IIx) were quantified in gastrocnemius muscle. Sections were air-dried for 30 minutes and then rehydrated with PBS for 5 minutes. Sections were then blocked with 0.5% BSA 0.5% Triton X-100 in PBS for 30 minutes at RT, then incubated with a cocktail of primary antibodies purchased from Developmental Studies Hybridoma Bank (DSHB, University of Iowa): BA-D5 (IgG2b, supernatant, 1:100 dilution) specific for MyHC-I, SC-71 (IgG1, supernatant, 1:100 dilution) specific for MyHC-IIa and BF-F3 (IgM, supernatant, 1:7.5 dilution) specific for MyHC-IIb for 1 hr at RT. After 3 washes with PBS (5 min each), the sections were incubated with secondary antibody cocktail (Invitrogen) to selectively bind to each primary antibody: goat anti-mouse IgG1 conjugated with Alexafluor488 (to bind to SC-71); goat anti-mouse IgG2b conjugated with Alexafluor350 (to bind to BA-D5); goat anti-mouse IgM conjugated with Alexafluor594 (to bind to BF-F3) for 1 hr in the dark at RT. After 3 washes with PBS (5 min each) and a brief rinse in water, the sections were mounted in 85% glycerol in PBS for imaging. Type IIa fibers will appear green, IIb as red, I as blue and the fibers that are not stained by these antibodies will appear black and are classified as type IIx.

### Digital image capture and analysis

Digital images were captured using a 20x objective in a Leica DM4000B microscope equipped with Retiga 4000R digital camera (QImaging Systems, Surrey, BC, Canada). Background correction was conducted for all images using an unstained adjacent area of the slide. Images were converted to 8-bit format and inverted. Fiber type was classified based on immunostaining for myosin isoform and individual fibers were outlined using a free hand tool and the cross-sectional area. Stain intensity of the cytochrome c oxidase and succinate dehydrogenase was quantified using ImageJ/Fiji software. Using a custom-generated algorithm, the blue stained areas for collagen were separated and highlighted from the rest of the image and % stained area was quantified (MIPAR software). Mitochondrial analysis was performed on cells in ImageJ by applying image deconvolution, background subtraction, adaptive binarization, and segmentation algorithms followed by particle analysis and morphology analysis with the ImageJ plugin MiNA to quantify intensity and mitochondrial branching.

### Cell culture and reagents

NIH3T3 fibroblasts were purchased from ATCC and cultured in DMEM supplemented with 10% bovine serum and 1% Penicillin/Streptomycin. The cells were treated with vehicle (DMSO in media) or AdipoRon (10µM or 50µM in media) for 10 minutes (AMPK phosphorylation) or 60 minutes (gene expression analysis). After treatment, cells were washed once with PBS and then lysed in Trizol or RIPA buffer depending on the experiment.

### JC-1 assay and Oxygen Consumption

NIH 3T3 cells (1×10^6^) were grown in DMEM supplemented with 10% bovine serum and 1% Penicillin/Streptomycin. Next day, the cells were treated with vehicle or AdipoRon (50µM) for 2 or 4 hrs. For JC-1 assay, the cells washed once with PBS and incubated with 2µg/ml JC-1 (Sigma) in media for 30 minutes at 37C. The cells were then trypsinized and equal number of cells was loaded on to 96 well plates and read at emission wavelength 530 and 590nm, where the ratio of 530/590 indexed mitochondrial membrane potential. For oxygen consumption assays, cells were trypsinized and equal numbers (4×10^5^) were re-suspended in respiration buffer and loaded on to the PreSens Oxoplates (Regensburg, Germany). Appropriate ambient and anoxic controls were included in the same plate.

### ATP luminescence assay

Relative ATP levels were quantified (ATPLite™assay (Perkin-Elmer, Waltham, MA; INFINITE M1000 PRO microplate reader (TECAN, Grodig, Austria)) in cells cultured as above and treated with vehicle or AdipoRon (50µM) for the prescribed times.

### Cell Proliferation Assay

NIH 3T3 cell proliferation was quantified according to the CyQUANT® Cell Proliferation Assay Kit (Molecular Probes, Inc. Eugene OR, USA) at four cell densities 1600 – 6700 cells/well. In brief, cells were seeded in microplate wells in growth medium at desired densities along with serial dilutions of cells for determination of a cell number standard curve. Cells were incubated for 24h after which culture medium was removed, and the number of cells quantified according to manufacturer instructions.

### Nonhuman Primate PBMC isolation and culture

Blood was collected from primates at the Wisconsin National Primate Research Center (WNPRC) with the approval of the Institutional Animal Care and Use Committee of the Office of the Vice Chancellor for Research and Graduate Education of the University of Wisconsin, Madison. PBMCs were isolated from 6ml of blood using SepMate-50 tubes (Stemcell Technologies, Cambridge, MA, USA). Isolated primate PBMC’s (1×10^6^) were cultured in 10cm plates in DMEM supplemented with 10% fetal bovine serum and 1% Penicillin/Streptomycin. Cells were treated with vehicle (DMSO in media) or AdipoRon (10µM or 50µM in media) for 5 minutes (AMPK phosphorylation) or 60 minutes (gene expression analysis). PBMCs were collected by centrifugation and the cell pellet washed and lysed in Trizol or RIPA buffer.

### Statistical analysis

Results are expressed as mean ± pooled SEM. For cell culture and animal experiments, data sets involving 2 groups were analyzed by unpaired two-tailed t-tests and for data sets with 3 groups one-way ANOVA followed by Bonferroni’s multiple comparison test with a cutoff of p<0.05.

## ACKNOWLEDGMENTS

The authors would like to acknowledge support from the Department for Veterans Affairs VA Merit BX003846, BX004031, NIH/NIA AG040178, AG056771, NIH training fellowships AG000213 (AES) and GM083252 (PRH), and the Glenn Foundation for Medical Research. This publication was made possible in part by NIH/ORIP grant P51OD011106 to the Wisconsin National Primate Research Center, University of Wisconsin-Madison. This work was supported by the use of facilities and resources at the William S. Middleton Memorial Veterans Hospital Madison Wisconsin.

## Supplemental materials

**Table S1.**
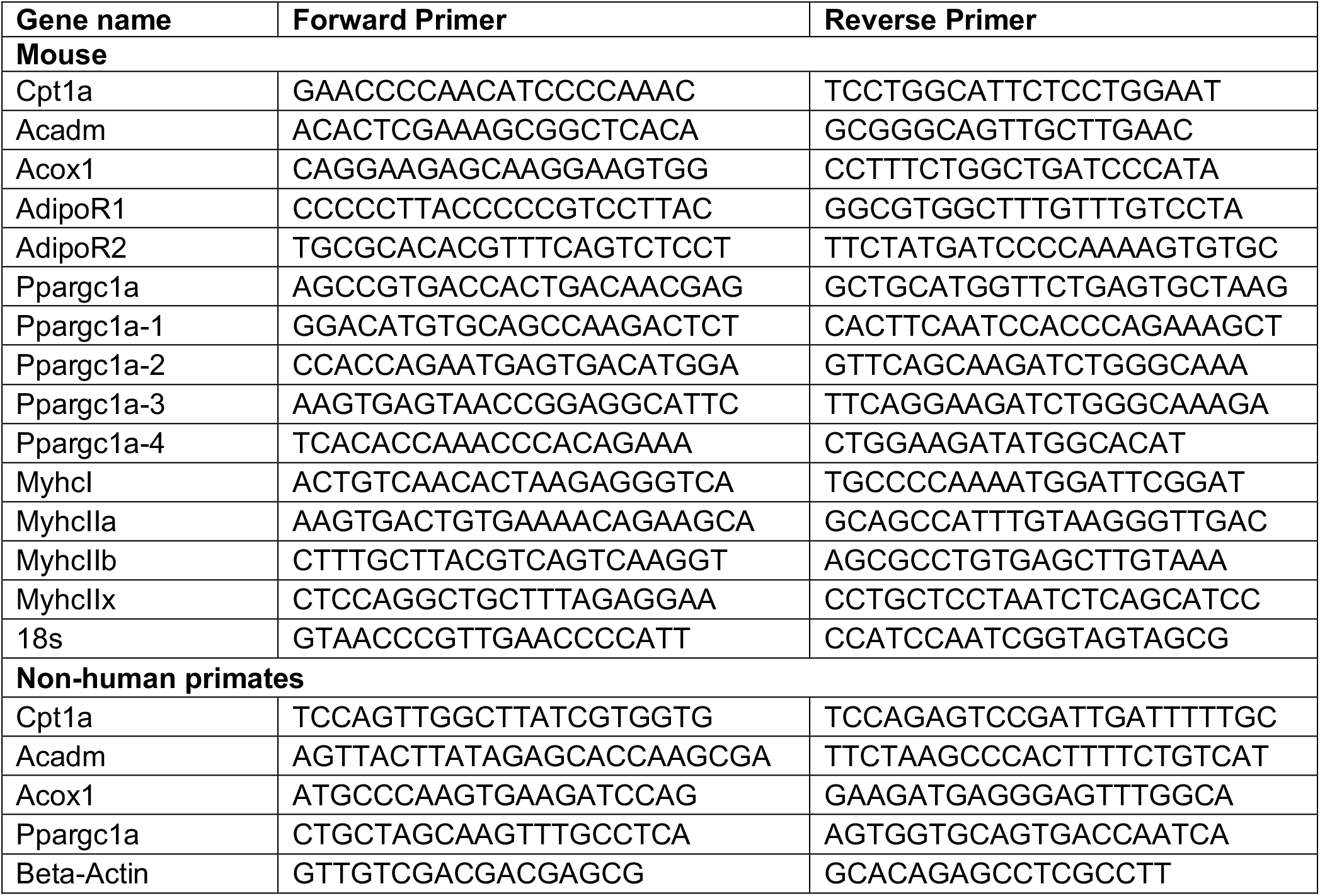
List of primers used for real time PCR analysis

**Figure S1:**
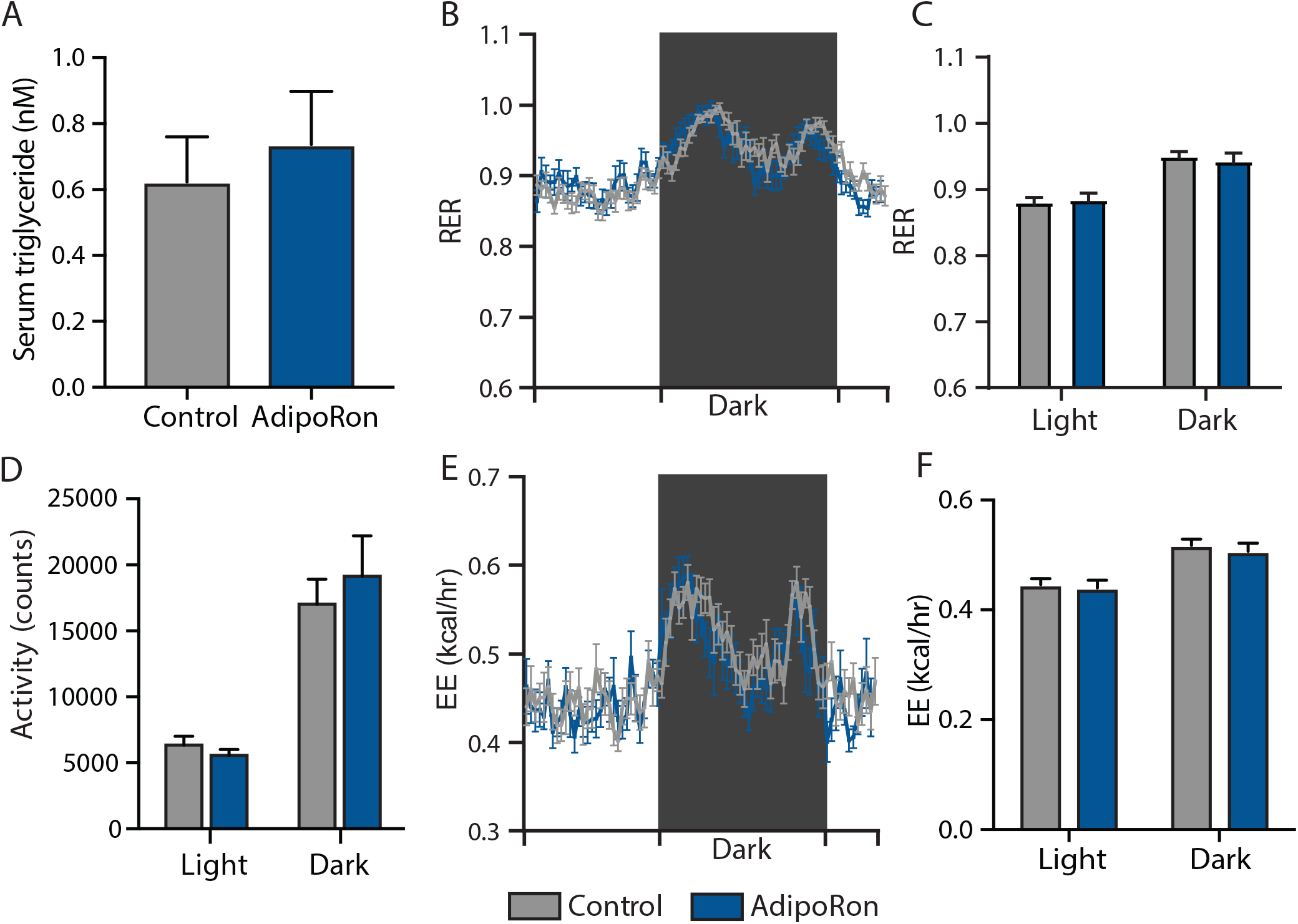
Effects of chronic AdipoRon treatment in aged mice. Effects of chronic AdipoRon treatment (1.2mg/kg BW, intravenous injection three times per week for 6 weeks) in aged male mice (25 months old). The following measures were conducted at the end of treatment period: (A) serum triglyceride levels. In metabolic chambers: (B) 24h respiratory exchange ration and light & dark mean (C), box signifies night (D) activity, (E) 24h energy expenditure, light & dark mean (F). Control (n=10) and AdipoRon (n=8). Data shown as average ±SEM.

**Figure S2:**
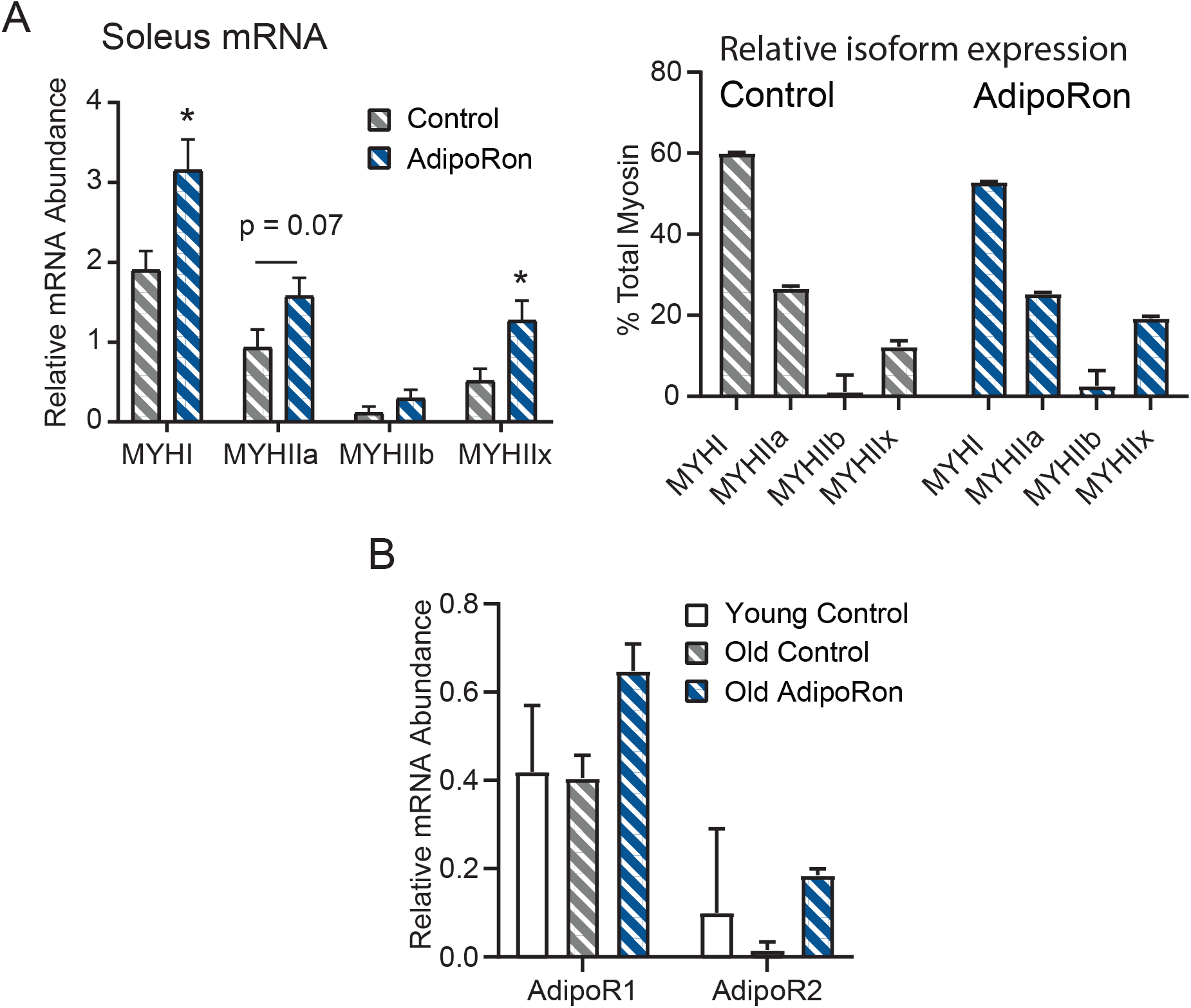
Chronic AdipoRon treatment in aged mice soleus. (A) Relative mRNA expression and percent isoform makeup of myosin heavy chain isoforms in aged control and AdipoRon treated mice. (B) mRNA expression effect of AdipoRon treatment in adiponectin receptors in young and aged control mice. Control (n=7-10) and AdipoRon (n=6-8). Data shown as arbitrary log-fold abundance units relative to 18s reference with 95% confidence interval determined via pooled SEM estimation and significance determined by Student’s t test (* p<0.05).

**Figure S3:**
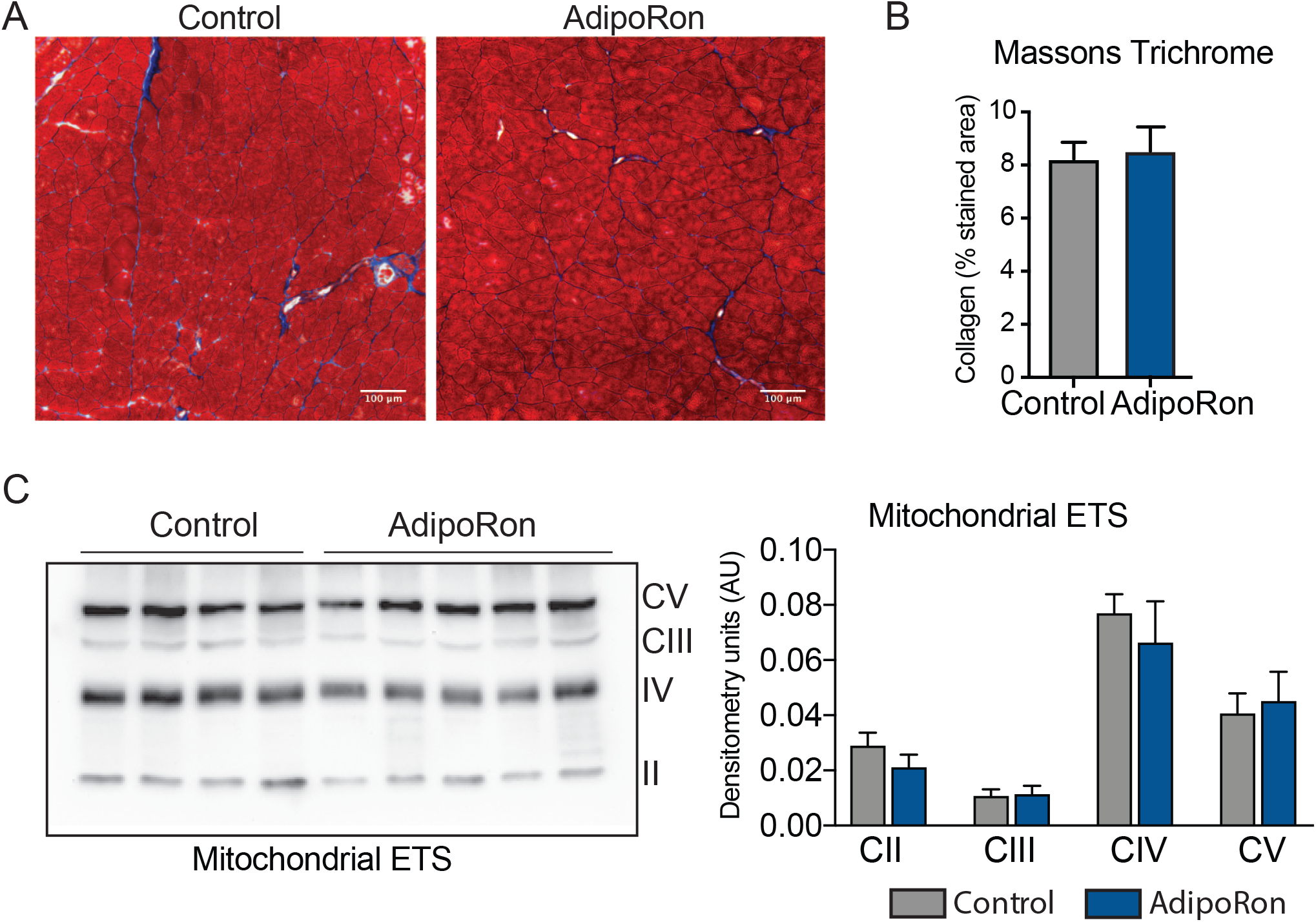
Effect of AdipoRon treatment on aged muscle fibrosis. Chronic AdipoRon treat-ment effect on muscle fibrosis shown via (A) representative Masson’s trichrome stained images from control and AdipoRon gastrocnemius, (B) Quantitation of intramuscular colla-gen content based on n=5 images per mouse, (C) Immunodetection of proteins in the electron transport system (ETS) in gastrocnemuis from control and AdipoRon treated animals with immunoblot quantitation relative to ponceau total protein loading control.

**Figure S4:**
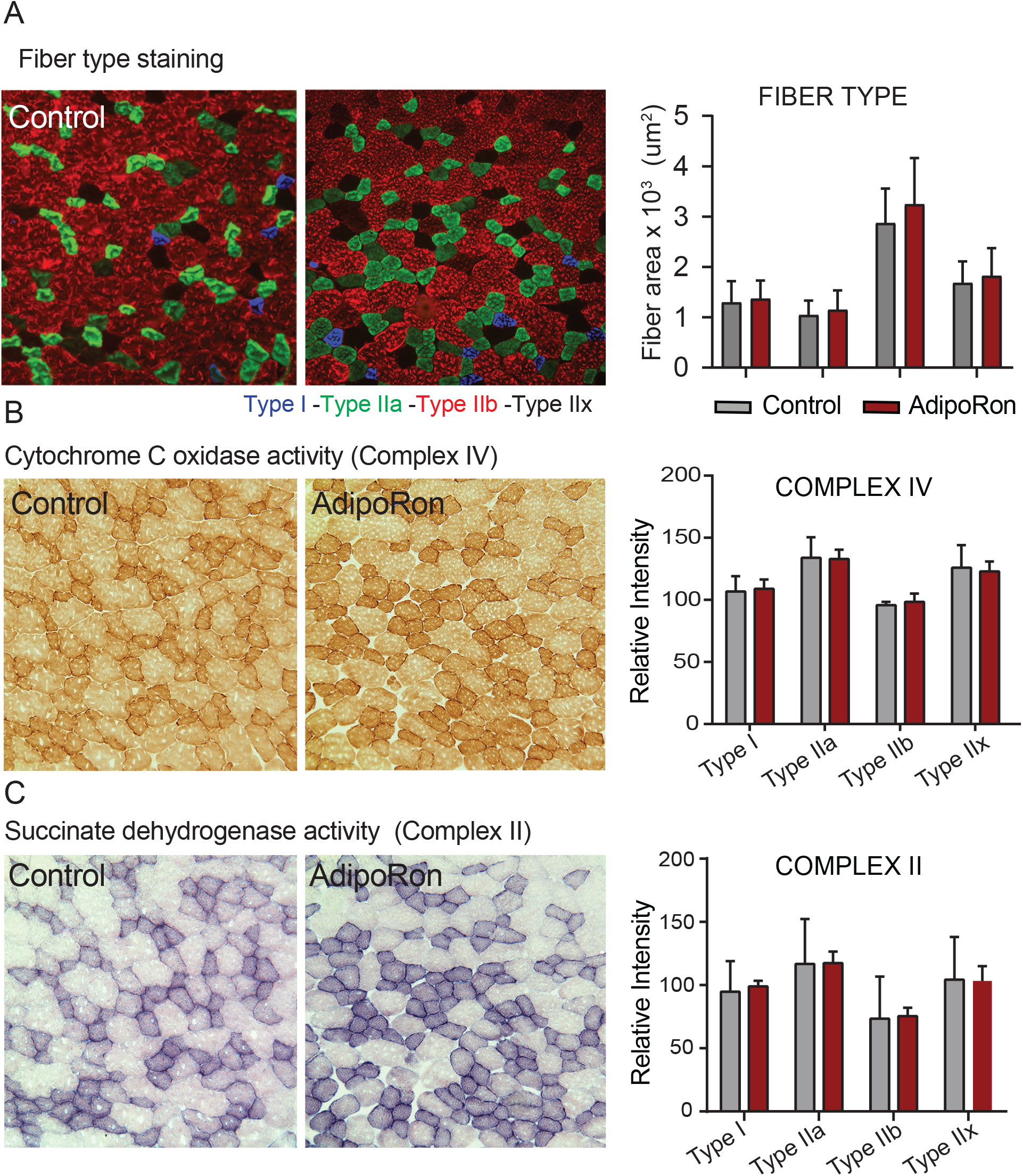
Effects of chronic AdipoRon treatment in young adult mice. Effects of chronic AdipoRon treatment (1.2mg/kg BW, intravenous injection three times per week for 6 weeks) in young B6C3F1 male mice (8-week-old). (A) Representative images of fiber typing immunohistochemistry for Type I (blue), Type IIa (green), Type IIb (red), and Type IIx (black) fibers in adjacent gastrocnemius muscle sections of control and AdipoRon treated animals. (B) Histochemical staining for maximal Cytochrome C Oxidase activity (Complex IV), (C) Histochemical staining for maximal succinate dehydrogenase (Complex II, middle panel), and their corresponding. Control (n=4) and AdipoRon (n=4). All data shown as average ±SEM.

**Figure S5:**
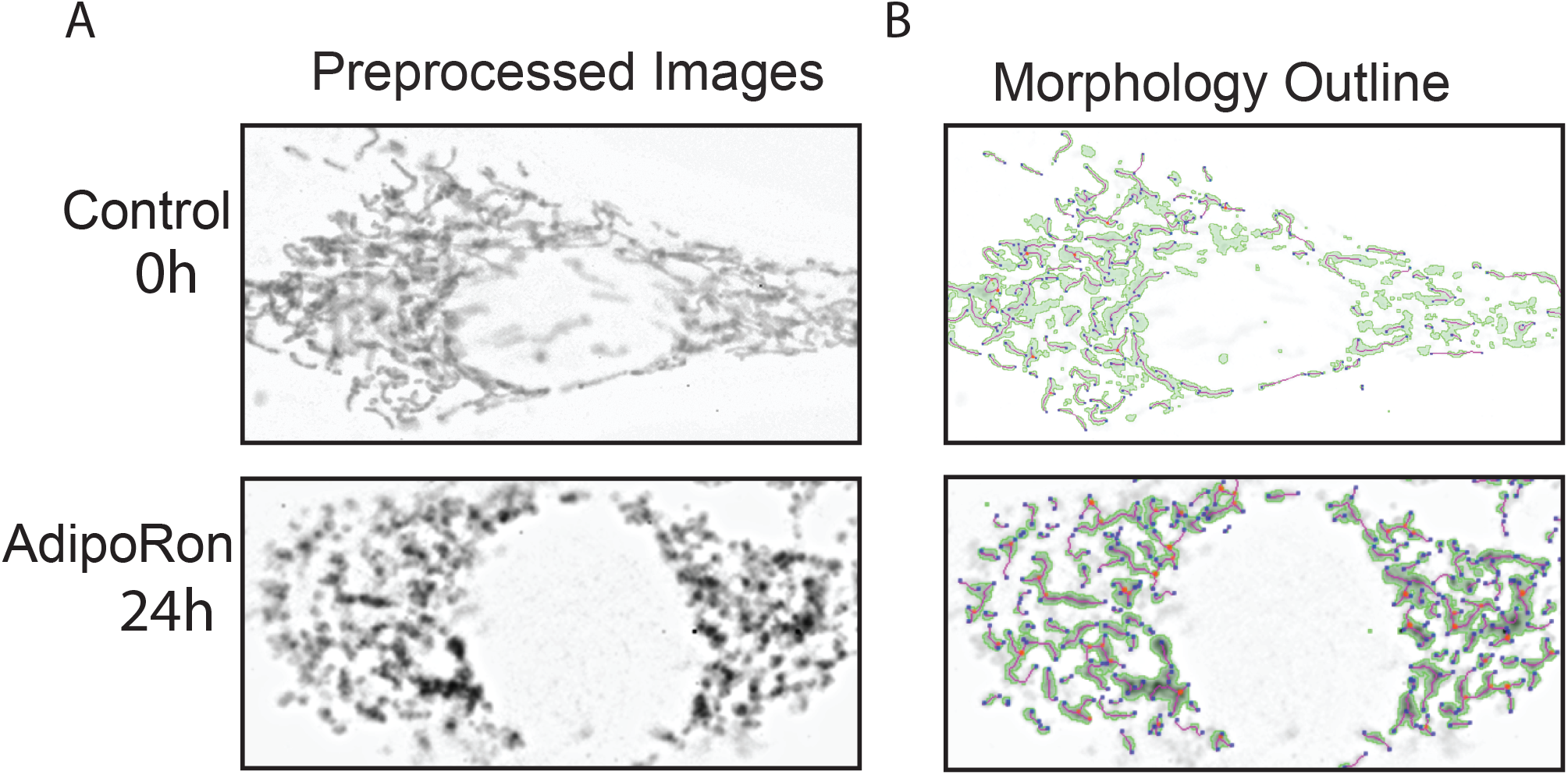
Mitochondrial Morphology Analysis. (A) Representative images of culture control and AdipoRon 50 um treated NIH-3T3 fibroblasts shown with intensity and contrast pre-processing filters applied, (B) Images after mitochondrial footprint morphology mask and 2D skeleton identified via MINA ImageJ plugin binarization and ridge detection algorithms.

